# An essential checkpoint for TLR9 signaling is release from Unc93b1 in endosomes

**DOI:** 10.1101/410092

**Authors:** Olivia Majer, Brian J Woo, Bo Liu, Erik Van Dis, Gregory M Barton

## Abstract

Nucleic acid-sensing Toll-like receptors (TLRs) are subject to complex regulation to facilitate recognition of microbial DNA and RNA while limiting recognition of self-nucleic acids^1^. Failure to properly regulate nucleic acid-sensing TLRs can lead to autoimmune and autoinflammatory disease^2-6^. Intracellular localization of these receptors is thought to be critical for self vs. non-self discrimination^7^, yet the molecular mechanisms that reinforce compartmentalized activation of intracellular TLRs remain poorly understood. Here we describe a new mechanism that prevents TLR9 activation from locations other than endosomes. This control is achieved through the regulated release of TLR9 from its trafficking chaperone Unc93b1, which only occurs within endosomes and is required for ligand binding and signal transduction. Mutations in Unc93b1 that increase affinity for TLR9 impair release and result in defective signaling. The release is specific to TLR9, as TLR7 does not dissociate from Unc93b1 in endosomes. This work defines a novel checkpoint that reinforces self vs. non-self discrimination by TLR9 and provides a mechanism by which TLR9 and TLR7 activation can be distinctly regulated.

## Main Text

To limit responses to extracellular self-nucleic acids, TLRs localize to intracellular compartments and require proteolytic cleavage within their ectodomains for activation^7-10^. The multi-pass transmembrane protein Unc93b1 is a trafficking chaperone for nucleic acid-sensing TLRs, facilitating trafficking of receptors from the ER to endosomes^11^. Unc93b1 is absolutely essential for endosomal TLR function, as humans and mice lacking functional Unc93b1 exhibit loss of endosomal TLR signaling and associated immunodeficiency^12,13^. Moreover, aberrant trafficking of endosomal TLRs due to mutations in Unc93b1 can lead to the breakdown of self versus non-self discrimination and contribute to autoimmune and autoinflammatory diseases in mouse models^3,14^. It remains unclear whether these alterations in TLR function are exclusively attributable to alterations in ER export. It has been suggested that Unc93b1 may regulate additional sorting steps to endosomes that are distinct between individual TLRs^15^, but recent work has argued that TLR function is unaltered when Unc93b1 is forcibly retained in the ER^16^. Thus, the mechanisms by which Unc93b1 contributes to the proper compartmentalization of endosomal TLRs and influences responses to different sources of nucleic acids remain unclear.

To dissect the mechanisms by which Unc93b1 regulates TLR signaling, we performed an unbiased alanine-scanning mutagenesis screen of Unc93b1. We stably expressed 204 individual FLAG-tagged Unc93b1 mutant alleles in a RAW264.7 macrophage cell line in which endogenous Unc93b1 was deleted by CRISPR/Cas9-mediated genome editing. To assess the impact of individual mutations on TLR function, we measured responses of each stable line after stimulation with ligands for Unc93b1-dependent TLRs (polyI:C for TLR3, R848 for TLR7, and CpG oligonucleotides for TLR9) and for an Unc93b1-independent TLR (LPS for TLR4). The non-functional allele Unc^H412R^ (HR), previously identified by Beutler’s group via an ENU mutagenesis screen^13^, served as a negative control.

The screen identified an Unc93b1 mutation (SKN to AAA at residues 282-284 in loop 5, hereafter referred to as Unc93b1^SKN^) that specifically disrupted signaling of TLR9, but not signaling by TLR7 or TLR3 (Figs. 1a,b). We first considered whether the mutation prevented TLR9 exit from the ER by introducing the Unc93b1^SKN^ mutant into Unc93b1-deficient RAW cell lines expressing an HA-tagged version of TLR9. Surprisingly, TLR9 trafficking to endosomes appeared normal, as the level of receptor cleavage, which occurs in endosomes, was similar between Unc93b1^WT^- and Unc93b1^SKN^-expressing cells (Fig. 1c). We confirmed trafficking of Unc93b1^SKN^ to Lamp1^+^ endosomes by immunofluorescence microscopy (Fig. S1a). If anything, the overall amount of Unc93b1 in endosomes seemed to be slightly higher in Unc93b1^SKN^-expressing cells compared to wildtype. Because the only described function for Unc93b1 is as a trafficking chaperone, a mutant version disrupting TLR9 signaling without abrogating trafficking to endosomes was unexpected and suggested we may have uncovered a new trafficking-independent function for Unc93b1.

**Fig. 1.**
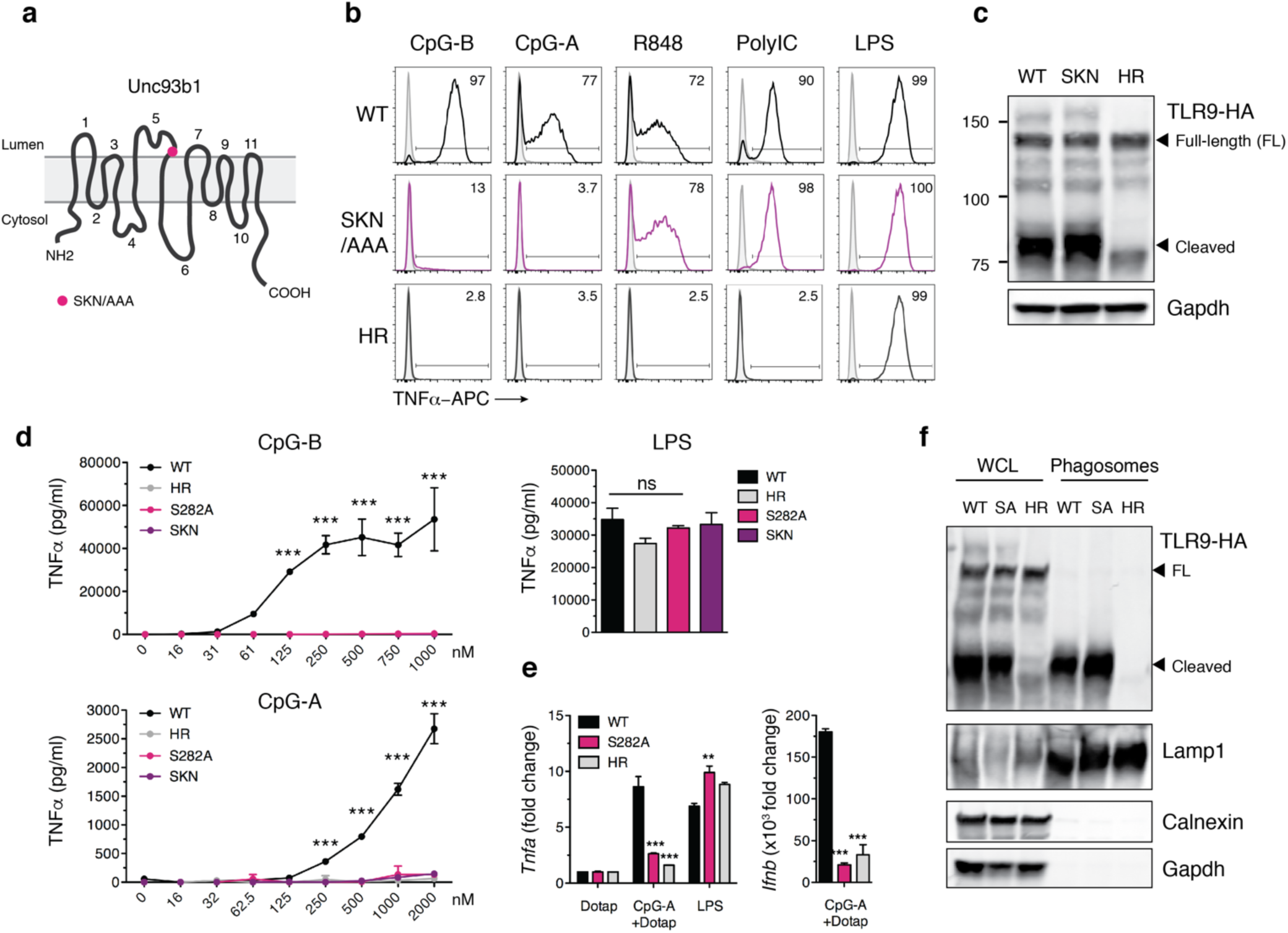
An Unc93b1 mutation in loop 5 results in defective TLR9 signaling despite normal trafficking. (**a**) Domain structure of Unc93b1 with position of the SKN mutation. (**b**) Unc93b1^SKN^ selectively impairs TLR9 signaling in RAW macrophages. Representative flow cytometry analysis showing percent TNFα positive cells, measured by intracellular cytokine staining of Unc93b1-deficient RAW macrophages retrovirally transduced to express the indicated Unc93b1 alleles (WT, SKN, and non-functional HR) after stimulation with CpG-B and CpG-A (150nM) for TLR9, R848 (25ng/ml) for TLR7, PolyIC (500ng/ml) for TLR3, or LPS (10ng/ml) for TLR4. For PolyIC responses RAW macrophages were additionally retrovirally transduced to express TLR3-HA. (**c**) TLR9 trafficking is normal in Unc93b1^SKN^ macrophages. Immunoblot of TLR9 from lysates of RAW macrophage lines shown in (**b**). (**d**) Unc93b1^S282A^ is sufficient for the TLR9 signaling defect. TNFα production, measured by ELISA, from the indicated RAW macrophage lines after stimulation for 8h with CpG-B, CpG-A, or LPS (50ng/ml) (n=3, representative experiment of two independent repeats). (**e**) Quantitative RT-PCR analysis of *Tnfa* and *Ifnb* expression in the indicated RAW macrophage lines 8h after stimulation with CpG-A/DOTAP (1μM) or LPS (10ng/ml) (n=3, representative experiment of two independent repeats). (**f**) Unc93b1^S282A^ does not affect TLR9 trafficking to endosomes. Whole cell lysates (WCL) or lysates of phagosomes isolated from the indicated RAW macrophage lines were probed for levels of TLR9, Lamp1, Calnexin, and GAPDH by immunoblot; FL: full-length. All data are mean ± SD; *P < 0.05, **P < 0.01, ***P < 0.001 by unpaired Student’s t-test (qRT-PCR) or two-way ANOVA followed by a Bonferroni posttest (ELISA). The data are representative of at least three independent experiments, unless otherwise noted.

To validate these findings in primary cells, we transduced bone marrow-derived macrophages (BMMs) from *Tlr9*^HA:GFP^*Unc93b1*^-/-^ mice with a retroviral vector driving expression of Unc93b1^SKN^ and tested TLR9 signaling. In *Tlr9*^HA:GFP^ mice the endogenous TLR9 gene has been modified to encode a C-terminal HA tag^17^. Again, TLR9 signaling was completely absent in Unc93b1^SKN^ BMMs despite normal receptor trafficking and a complete restoration of TLR7 signaling (Figs. S1b,c). To determine if a mutation of any single amino acid within the SKN motif was sufficient to recapitulate loss of TLR9 signaling, we individually mutated each amino acid within the SKN motif to alanine. We found that the single S282A (SA) mutation was sufficient to abolish TLR9 signaling in an NF-*k*B luciferase reporter assay in HEK293T cells, whereas K283A and N284A showed no effect (Fig. S1d). Similar to the Unc93b1^SKN^ mutant, Unc93b1^S282A^-expressing RAW macrophages failed to respond to CpG-B or CpG-A oligonucleotides, while TLR7 and TLR3 responses were unaffected (Fig. 1d and Fig. S1e). TLR9-dependent induction of IFNß after stimulation with CpG-A/Dotap was also abrogated (Fig. 1e). Again, the level of cleaved TLR9 in phagosomes was similar compared to wildtype, indicating normal trafficking (Fig. 1f). These results indicate that mutation of Ser282 in Unc93b1 is sufficient to abrogate TLR9 activation without altering TLR9 trafficking or influencing other Unc93b1-dependent TLRs.

We next investigated how Unc93b1^S282A^ interferes with TLR9 signaling. Based on the luminal position of the mutation (Fig. 1a), we reasoned that ligand recognition could be affected rather than signal transduction after ligand binding. To assess ligand binding, we fed RAW macrophages biotin-conjugated CpG-B, pulled down the ligand with streptavidin-beads, and compared the amount of bound TLR9. Unc93b1^S282A^ mutant cells showed significantly less TLR9 bound to CpG-B, suggesting a reduced affinity of TLR9 for ligand (Fig. 2a). To rule out differences in DNA uptake or sampling, we verified that all cell lines were equally capable of endocytosing CpG-B (Fig. 2b) and that the ligands were effectively delivered to Lamp1^+^ endolysosomes containing TLR9 (Fig. S2).

**Fig. 2.**
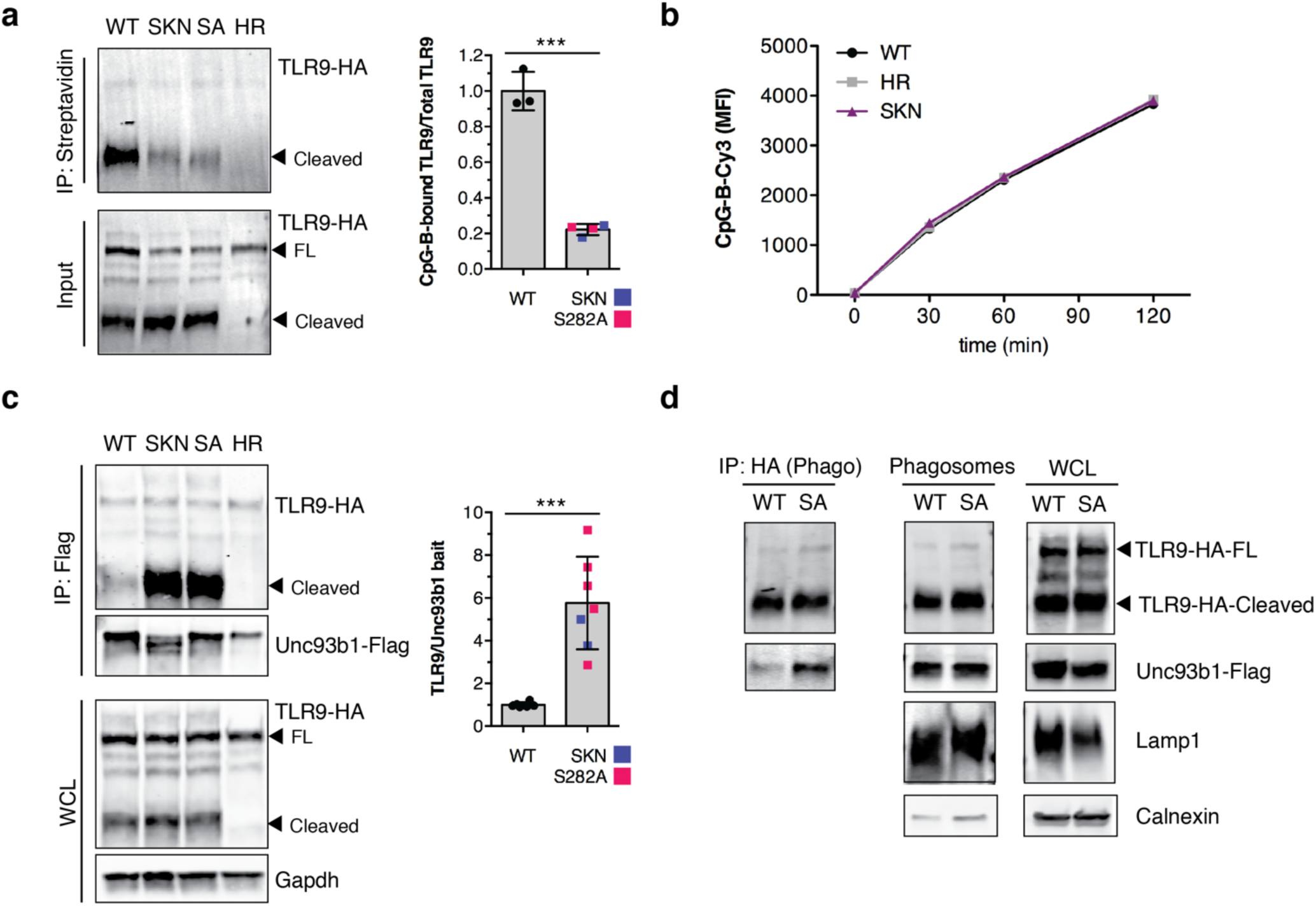
Unc93b1^S282A^ attenuates ligand binding and increases association with TLR9. (**a**) Unc93b1^SKN^ and Unc93b1^S282A^ attenuate ligand binding of TLR9. Streptavidin immunoprecipitation of lysates from RAW macrophage lines, expressing TLR9-HA and the indicated Unc93b1 alleles, stimulated for 4h with biotinylated CpG-B (1μM) followed by immunoblot for TLR9. Bar graph shows quantification of CpG-bound TLR9 over total TLR9 from several experiments. (**b**) Unc93b1^SKN^ cells show normal DNA uptake. Uptake of Cy3-labeled CpG-B (1μM) as measured by flow cytometry of RAW macrophage lines expressing the indicated alleles of Unc93b1. (**c**) Unc93b1^SKN^ and Unc93b1^S282A^ bind stronger to TLR9. Flag immunoprecipitation of Unc93b1 from RAW macrophage lines, expressing TLR9-HA and the indicated Unc93b1 alleles, followed by immunoblot of TLR9. Input levels of TLR9 in whole cell lysates (WCL) are also shown. Bar graph shows quantification of TLR9 bound to Unc93b1 from several experiments. (**d**) HA immunoprecipitation of TLR9 from phagosome preparations of RAW macrophage lines, expressing TLR9-HA and the indicated Unc93b1 alleles, followed by immunoblot for Unc93b1. Input levels of TLR9 and Unc93b1 in phagosomes and whole cell lysates (WCL) are also shown; FL: full-length. Representative of two independent experiments. All data are mean ± SD; *P < 0.05, **P < 0.01, ***P < 0.001 by unpaired Student’s t-test. The data are representative of at least three independent experiments, unless otherwise noted.

We considered the possibility that a loss of interaction between Unc93b1^S282A^ and TLR9 during trafficking could impair ligand binding and activation of TLR9. On the contrary, we observed greater interaction between TLR9 and the mutants Unc93b1^SKN^ and Unc93b1^S282A^ when compared to wildtype Unc93b1 (Figs. 2c and S3a). TLR7, an endosomal TLR whose signaling is unaffected by Unc93b1^S282A^, did not show greater interaction with either Unc93b1 mutant (Fig. S3b). Furthermore, stronger association with TLR9 was only observed for the S282A mutation, but not for mutations of the neighboring amino acids K283 and N284 (Fig. S3c). We confirmed the increased interaction between TLR9 and Unc93b1^S282A^ in endosomes directly by isolating phagosomes with magnetic beads and measuring TLR9-Unc93b1 association by immunoprecipitation. Again, a higher interaction between TLR9 and Unc93b1^S282A^ was observed (Fig. 2d). Taken together, these results show that TLR9 signaling inversely correlates with the extent of interaction between TLR9 and Unc93b1 and suggest that a stronger association between Unc93b1 and TLR9 may interfere with ligand binding by TLR9.

To better understand how the S282A mutation affects the association between Unc93b1 and TLR9, we examined the importance of neighboring residues within loop 5 of Unc93b1. We analyzed all triple-alanine mutants within our mutant library that spanned loop 5 of Unc93b1 for their impact on TLR9 signaling and binding (Fig. 3a). Scattered throughout loop 5 were several Unc93b1 mutants that affected TLR9 signaling. However, within those identified mutations, five of them occurred in a row (spanning amino acids 270-284 and containing Ser282) and showed impaired TLR9 signaling, suggesting the existence of a larger region important for TLR9 function (Fig. 3b). When probing Unc93b1 mutants within this region for their impact on TLR9 binding, we found that mutations in the core part of this region (amino acids 276-281) completely abolished the interaction with TLR9 (Fig. 3c), whereas mutations in the boundaries of the region (amino acids 273-275 and 282-284), including Ser282, increased the interaction. Notably, both loss and increase of the Unc93b1-TLR9 interaction resulted in impaired TLR9 signaling, highlighting the importance of an optimal binding affinity for proper TLR9 function. Note that mutation of amino acids 279-281 resulted in constitutive activation (Fig. 3b). We suspect that this TNFα production in the absence of TLR stimulation may be due to ER stress stemming from the abnormally high expression level of this Unc93b1 mutant (Fig. 3c). Despite constitutive levels of TNFα, R848 and LPS were able to further induce cytokine levels whereas CpG-B was not, suggesting an impaired TLR9 response also for this mutant. Most of the mutations had little or no effect on TLR7 function (Fig 3b). Thus, these analyses have identified a region of Unc93b1 that is critical for interaction with TLR9 and demonstrate that differential regulation of Unc93b1-dependent TLRs can be mediated through unique interactions between Unc93b1 and each receptor.

**Fig. 3.**
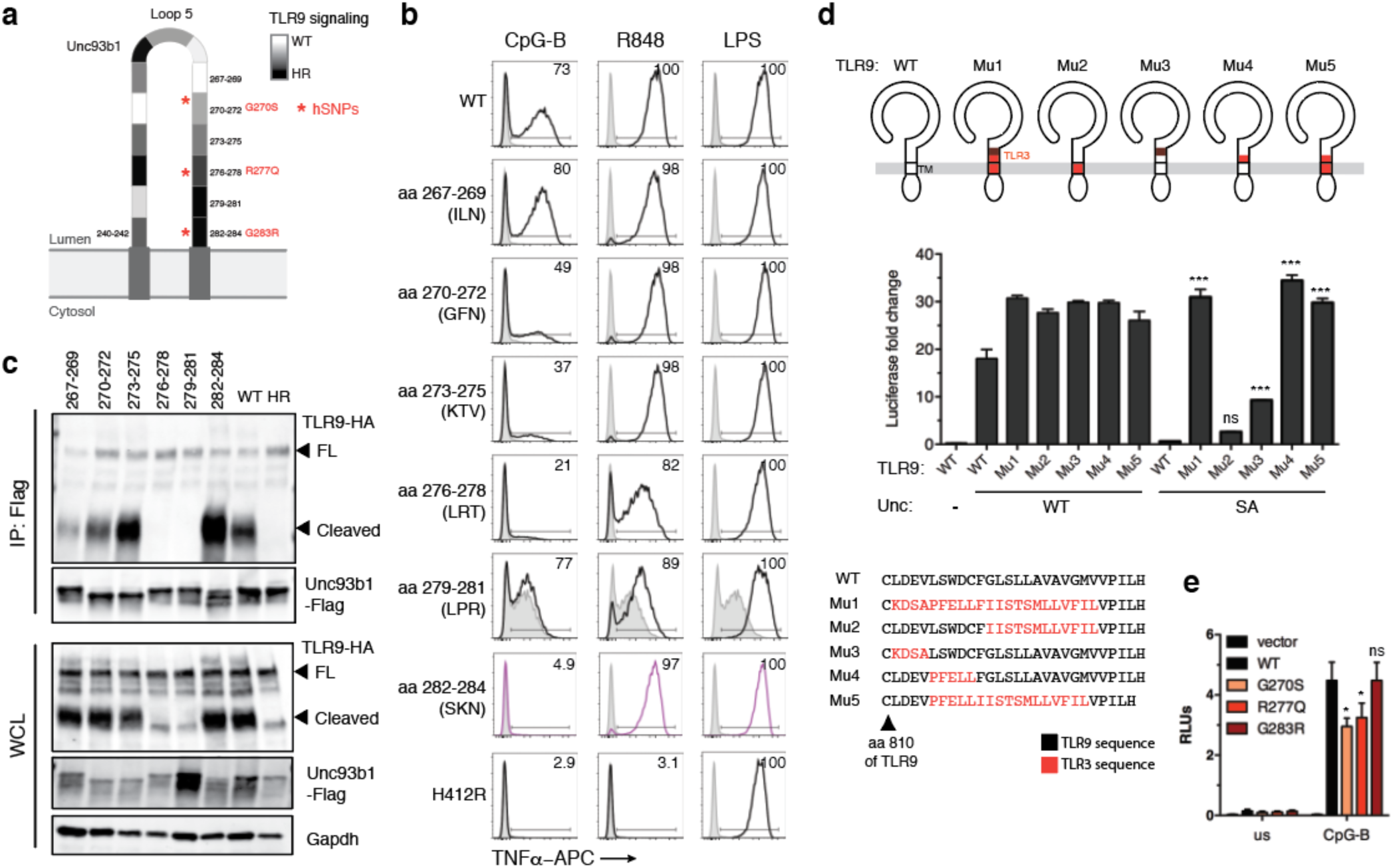
Identification of residues within loop 5 of Unc93b1 that mediate interaction with TLR9. (**a**) Schematic of the tested mutants in loop 5 of Unc93b1 and the relative TLR9 responses indicated in shades of grey: white indicates a response equivalent to WT while black indicates no response. (**b**) A larger region in loop 5 of Unc93b1 is important for TLR9 signaling. Representative flow cytometry analysis showing percent TNFα positive cells, measured by intracellular cytokine staining of RAW macrophages retrovirally transduced to express the indicated Unc93b1 mutants (spanning amino acids 267-284, and non-functional HR) after stimulation with CpG-B (25nM) for TLR9, R848 (100ng/ml) for TLR7, and LPS (10ng/ml) for TLR4. Representative of three independent experiments. (**c**) A larger region in loop 5 of Unc93b1 mediates binding to TLR9. Flag immunoprecipitation of Unc93b1 from RAW macrophage lines, expressing TLR9-HA and the indicated Unc93b1 alleles, followed by immunoblot of TLR9. Input levels of TLR9 and Unc93b1 in whole cell lysates (WCL) are also shown; FL: full-length. Representative of two independent experiments. (**d**) Schematics showing relative positions and a sequence alignment (bottom) of swapped regions within the TLR9 chimeras. NF?B activation was measured by luciferase assay in HEK293T cells transiently transfected with the indicated TLR9 and Unc93b1 variants and stimulated with CpG-B (200nM) for 16h. Data are normalized to Unc93b1-independent hIL-1b responses and expressed as luciferase fold change over unstimulated controls (n=4, representative of two independent experiments). One-way ANOVA results: F(5/12)=300.0, *p*<0.0001 (**e**) NF-?B luciferase assay in HEK293T cells expressing TLR9 and the indicated human Unc93b1 variants and stimulated with CpG-B (250nM) for 16hrs. Data is normalized to Renilla expression and expressed as relative luciferase units (RLUs) (n=7, pooled from two independent experiments). One-way ANOVA results: F(3/12)=9.45, *p*=0.0017 All data are mean ± SD; *P < 0.05, **P < 0.01, ***P < 0.001 by one-way ANOVA followed by a Tukey’s posttest.

To identify the reciprocal binding region in TLR9 that engages with the loop 5 region of Unc93b1, we turned to a set of TLR9 chimeric variants in which varying segments of the juxtamembrane and transmembrane domains had been replaced with the corresponding sequence of TLR3^4^. As TLR3 is unaffected by Unc93b1^S282A^ (Fig. S1d) we reasoned that this strategy could identify critical residues within TLR9 that mediate interaction with Unc93b1. We compared signaling of each TLR9 variant in cells expressing Unc93b1^WT^ or Unc93b1^S282A^. These analyses identified a short motif of five amino acids (LSWDC) in the juxtamembrane region of TLR9 that when reverted to the corresponding TLR3 sequence (PFELL) restored function in the presence of Unc93b1^S282A^ (Fig. 3d). We conclude that this short motif within TLR9 engages with loop 5 of Unc93b1; alterations in the residues of these interaction surfaces can increase or decrease TLR9-Unc93b1 binding, which influences TLR9 ligand binding and function.

The amino acids of loop 5 implicated in interaction with TLR9 are completely conserved between mouse and human, so we sought to validate the functional importance of this region for human Unc93b1. To do so, we looked for naturally occurring human variants in residues predicted to influence TLR9 binding. Three SNPs have been reported in the NCBI Single Nucleotide Polymorphism Database (Fig. 3a), all occurring with very low frequencies (Minor Allele Frequencies between 8.5 - 9.1E-06). We cloned the three human Unc93b1 variants and tested them in HEK293T cells for their effect on TLR9 signaling. Two variants (G270R and R277Q) positioned within the critical Unc93b1 binding region significantly reduced TLR9 signaling, whereas G283S had no effect (Fig. 3e). The reduction in signaling was specific to TLR9, as none of the human Unc93b1 variants reduced TLR7 signaling (Fig. S4). These results suggest that human genetic variants in this region of Unc93b1 may affect TLR9 function.

Based on the finding that an enhanced association with Unc93b1 is detrimental to TLR9 function, we reasoned that TLR9 might require release from Unc93b1 in endosomes prior to ligand binding and activation. If this model is correct, then (1) the association between TLR9 and Unc93b1 should be stronger in the ER and accordingly decrease in endosomes and (2) Unc93b1 should be absent from the active signaling complex of TLR9. We investigated each of these predictions in turn. First, we used cellular fractionation to separate ER and endosomes and measured the extent of TLR9-Unc93b1 interaction in each organelle preparation (Fig. 4a). We immunoprecipitated TLR9 from pooled fractions enriched for endosomes or ER and measured the amount of associated Unc93b1 (Fig. 4a). Consistent with a release of TLR9 from Unc93b1 in endosomes, the association of Unc93b1 and TLR9 was greater in the ER than in endosomes (Fig. 4b). Furthermore, Unc93b1^S282A^ showed an overall stronger association with TLR9, both in the ER and in endosomes, suggesting that the altered chemistry of the mutant might have an overall “sticky” effect on TLR9 that prevents efficient release in endosomes. Next, to test whether Unc93b1 is absent from the active TLR9 signaling complex, we immunoprecipitated the signaling adaptor MyD88 after stimulation of cells and probed for associated TLR9 and Unc93b1. We were able to pull down abundant amounts of cleaved TLR9 in wildtype cells, as previously reported^8^; however, Unc93b1 was not detectable within the signaling complex (Fig. 4c). As expected, Unc93b1^S282A^-expressing cells did not recruit MyD88 after stimulation, in accordance with their defective TLR9 response.

**Fig. 4.**
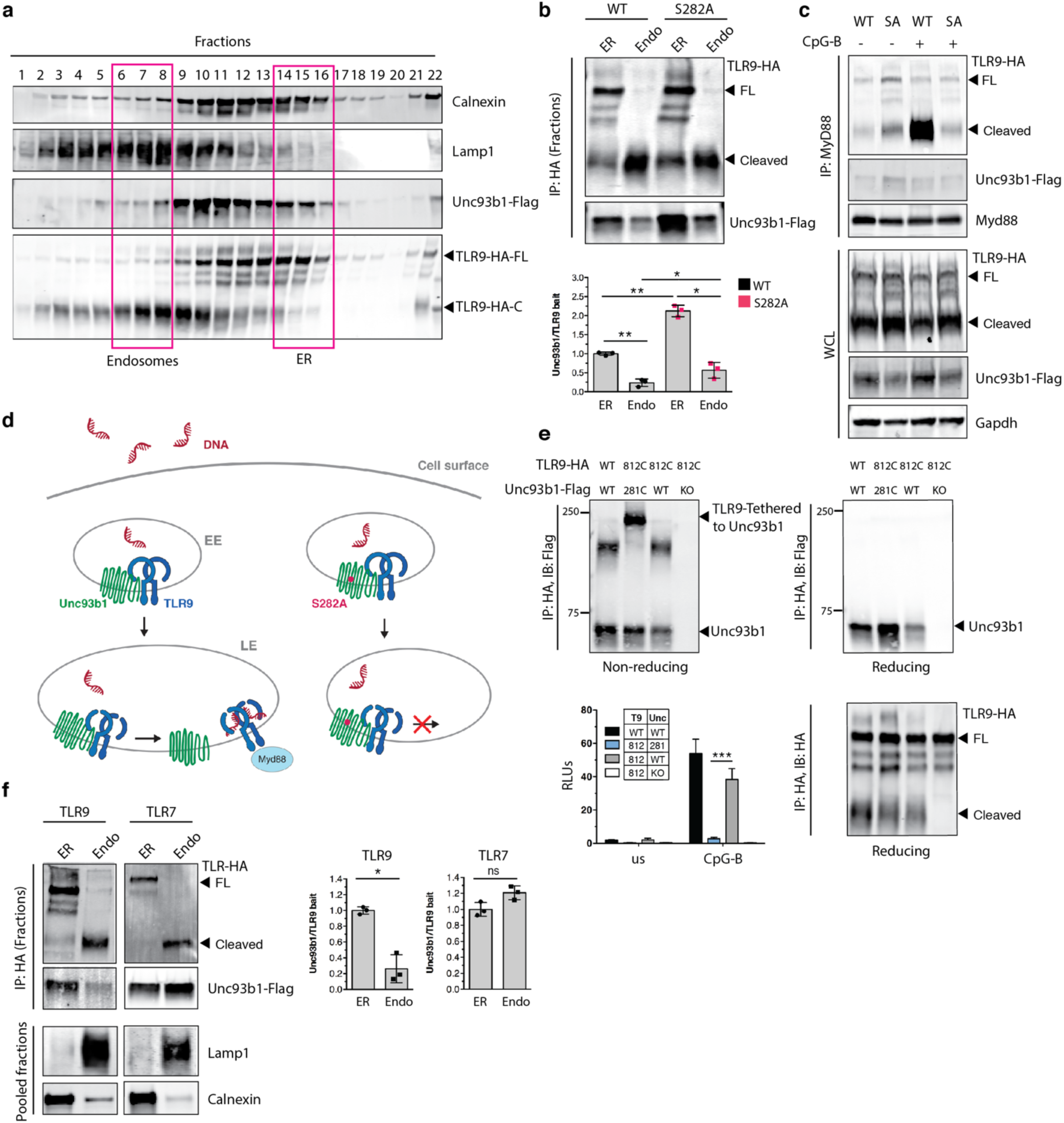
TLR9, but not TLR7, must release from Unc93b1 for signaling. (**a**) Sub-cellular fractionation of TLR9-HA, Unc93b1-FLAG expressing RAW macrophages was performed by density-gradient centrifugation. The distributions of Calnexin, Lamp1, Unc93b1, and TLR9 were measured by immunoblot. The pooled fractions, enriched for endosomes or ER, used for subsequent coimmunoprecipitation experiments are highlighted. (**b**) The TLR9-Unc93b1 association is reduced in endosomes compared to the ER. HA immunoprecipitation of TLR9 from pooled ER or endosome fractions (as shown in (**a**)) from RAW macrophage lines expressing TLR9-HA and the indicated Unc93b1 alleles. Immunoprecipitated TLR9-HA levels were normalized across fractions and probed for levels of Unc93b1. Bar graph shows the quantification of Unc93b1 bound to TLR9 across fractions. (**c**) Unc93b1 is absent from the active TLR9 signaling complex. Immunoprecipitation of MyD88 from RAW macrophage lines expressing TLR9-HA and the indicated Unc93b1 alleles and stimulated for 1h with CpG-B (1μM), followed by immunoblot for TLR9 and Unc93b1. Shown are also input levels of TLR9 and Unc93b1 in whole cell lysates (WCL). Representative of two independent experiments. (**d**) Release model of TLR9. (**e**) Unc93b1-tethered TLR9 is unable to signal. HA immunoprecipitation of TLR9 under non-reducing conditions from RAW macrophage lines expressing the indicated TLR9 and Unc93b1 cysteine mutants. Eluted proteins were either treated with the reducing agent dithiothreitol or left untreated and probed for Unc93b1. The non-reducing condition visualizes disulfide-bond formation between Unc93b1 and TLR9, which disappears after treatment with dithiothreitol (reducing condition). Bar graph shows an NF-?B luciferase assay in HEK293T cells expressing the indicated cysteine mutant combinations and stimulated with CpG-B (1μM) for 16hrs. Data are normalized to Renilla expression and expressed as relative luciferase units (RLUs) (n=4; representative of three independent experiments). (**f**) TLR7 does not release from Unc93b1 in endosomes. HA immunoprecipitation of TLR9 and TLR7 from pooled ER or endosome fractions of RAW macrophage lines expressing WT Unc93b1. Immunoprecipitated TLR-HA levels were normalized across fractions and probed for levels of Unc93b1. Bar graph shows the quantification of Unc93b1 bound to TLR9 or TLR7 between ER and endosome fractions; FL: full-length. All data are mean ± SD; *P < 0.05, **P < 0.01, ***P < 0.001 by paired Student’s t-test (**b**, **f**) or unpaired Student’s t-test (**e**). The data are representative of at least three independent experiments, unless otherwise noted.

Based on these results we propose a release model, whereby TLR9 must dissociate from Unc93b1 for efficient ligand binding to overcome the activation threshold for signaling. We speculate that Unc93b1 might interfere with DNA recognition by keeping the receptor in an unfavorable conformation for ligand binding, which would explain the attenuated ligand binding of TLR9 when forcefully bound to Unc93b1^S282A^ (Fig. 4d). To independently test this model, we sought to engineer a cysteine bridge between TLR9 and Unc93b1 to permanently tether the two proteins together and prevent release (Fig. S5a). We screened a panel of TLR9 and Unc93b1 cysteine mutants, focusing on the previously identified loop 5 binding region of Unc93b1 and the juxtamembrane region of TLR9. We identified a pair of Unc93b1 and TLR9 cysteine mutants (Unc93b1^281C^ and TLR9^812C^) that trafficked to endosomes (Fig. 4e, bottom right) yet remained attached through an intermolecular disulfide bond, which could be visualized by SDS-PAGE under non-reducing conditions (Fig. 4e, top left, and Fig. S5b). Preventing release from Unc93b1 (i.e., coexpression of Unc93b1^281C^ and TLR9^812C^) completely abrogated TLR9 signaling (Fig. 4e, bottom left). Importantly, Unc93b1^281C^ and TLR9^812C^ were both functional when expressed with wildtype TLR9 and Unc93b1, respectively, ruling out the possibility that the cysteine mutations simply created non-functional proteins (Fig. 4e, bottom left, and Fig. S5c).

Finally, we asked if TLR7 also requires release from Unc93b1 for signaling. We separated ER from endosomes and compared the association of TLR7 and Unc93b1. Surprisingly, the interaction did not decrease in endosomes (Fig. 4f), suggesting that, unlike TLR9, TLR7 can bind ligand and signal while associated with Unc93b1. In fact, in an accompanying manuscript^18^, we describe how the continued association of TLR7 and Unc93b1 in endosomes facilitates the regulation of TLR7 signaling through a distinct mechanism from the one we describe here for TLR9. Instead of inhibiting ligand binding, Unc93b1 recruits a negative regulator of TLR7 signaling. Disruption of this interaction leads to TLR7-dependent autoimmunity^18^. Thus, Unc93b1 utilizes distinct mechanisms to regulate activation of TLR9 and TLR7 in endosomes.

The differential regulation of TLR9 and TLR7 that we describe may explain the enigmatic observation of TLR9 and TLR7 contributing distinctly to the pathology of certain autoimmune diseases^19,20^. Inhibition of TLR9 function is strictly linked to proper trafficking, both through Unc93b1 association, as we describe here, and through the requirement for ectodomain proteolysis. Accordingly, overexpression of TLR9 does not induce disease^17^. In contrast, TLR7 appears subject to more ‘tunable’regulation that dampens but does not eliminate signaling, and overexpression of TLR7 is sufficient to break tolerance and drive autoimmunity^2,5,6^. Why distinct mechanisms of regulation have evolved for such functionally similar innate receptors remains unclear. One possibility is that differences in the trafficking of TLR7 and TLR9 influences the likelihood that self RNA or DNA will be encountered; indeed, TLR9 traffics to endosomes via the plasma membrane while TLR7 is thought to bypass the plasma membrane^15^. Alternatively, the nature of the ligands recognized by each receptor may require differing degrees of tunability. Recent work has revealed that TLR7 and TLR8 ligands are quite simple (e.g., TLR7 recognizes the purine nucleoside guanosine together with a 3-mer uridine-containing ssRNA)^21-23^. In this case, avoiding self-recognition may require more subtle modulation of signaling than is necessary for TLR9. Regardless of any teleological rationale, dissecting the mechanisms that underlie differential regulation of these TLRs should reveal new avenues for therapeutic manipulation of TLR activation.

## Methods

### Antibodies and Reagents

The following antibodies were used for immunoblots, immunoprecipitations, or flow cytometry: rat anti-HA as purified antibody or matrix (3F10; Roche), mouse anti-FLAG as purified antibody or matrix (M2; Sigma-Aldrich), anti-mLamp-1 (goat polyclonal, AF4320; R&D Systems), anti-calnexin (rabbit polyclonal; Enzo Life Sciences, ADI-SPA-860-F), anti-Myd88 (AF3109, R&D Systems), anti-Gapdh (GT239, GeneTex), anti-TNFα-APC (MP6-XT22; eBioscience), purified anti-CD16/32 Fc Block (2.4G2), goat anti-mouse IgG-AlexaFluor680 (Invitrogen), goat anti-mouse IgG-AlexaFluor680 (Invitrogen), rabbit anti-goat IgG-AlexaFluor680 (Invitrogen), goat anti-mouse IRDye 800CW (Licor), donkey anti-rabbit IRDye 680RD (Licor), goat anti-rat IRDye 800CW (Licor). For immunofluorescence: rat anti-HA (3F10; Roche), rabbit anti-Lamp1 (ab24170, Abcam), goat anti-rat AlexaFluor488 (Jackson Immunoresearch), goat anti-rabbit AlexaFluor647 (Jackson Immunoresearch). Cells were mounted in Vectashield Hard Set Mounting Medium for Fluorescence (Vector Laboratories). For ELISA: anti-mouse TNFa purified (1F3F3D4, eBioscience), anti-mouse TNFa Biotin (XT3/XT22, eBioscience), Streptavidin HRP (BD Pharmingen).

The following TLR ligands were used: CpG-B (ODN1668: TCCATGACGTTCCTGATGCT, all phosphorothioate linkages) and CpG-A (ODN1585: G*G*GGTCAACGTTGAG*G*G*G*G*G, asterix indicate phosphorothioate linkages) were synthesized by Integrated DNA Technologies (Cy3 or biotin was attached to the 5-prime end for imaging or biochemistry experiments), R848 (Invivogen), PolyIC HMW (Invivogen), and LPS (Invivogen). Human IL-1b was purchased from Invitrogen. NP-40 (Igepal CA-630) was purchased from Sigma-Aldrich. Saponin, was purchased from Acros. Lipofectamine-LTX reagent (Invitrogen) was used for transient transfection of plasmid DNA. DOTAP liposomal transfection reagent (Roche) was used for transfection of CpG-A. OptiMEM-I (Invitrogen) was used as media to form nucleic acid complexes for transient transfections. Streptavidin Magnetic Beads for biotin pull downs and Protein G agarose were purchased from Pierce. Violet fluorescent reactive dye for live dead staining of cells for flow cytometry was purchased from Invitrogen. ProMag 1 Series-COOH Surfactant free magnetic beads (#25029) for phagosome preparations were purchased from Polysciences. For renilla luciferase assays we used Coelenterazine native (Biotum). For firefly luciferase assays we used Luciferin (Biosynth). Assays were performed in Passive Lysis Buffer, 5x (Promega).

### Mice

Mice were housed under specific-pathogen-free conditions at the University of California, Berkeley. All mouse experiments were performed in accordance with the guidelines of the Animal Care and Use Committee at UC Berkeley. *Tlr9*^HA:GFP^ mice were generated as previously described^17^.

### Plasmid constructs

AccuPrime Pfx DNA polymerase (Invitrogen) was used for site directed mutagenesis using the QuikChange II Site-directed Mutagenesis protocol from Agilent Technologies. The following mouse stem cell virus (MSCV)-based retroviral vectors were used to express UNC93B1, TLR9, TLR7, and TLR3 in cell lines: MSCV-PuromCherry (IRES – PuromycinR-T2A-mCherry), MSCV2.2 (IRES-GFP), MSCV-Thy1.1 (IRES-Thy1.1), MIGR2 (IRES-hCD2). 3× FLAG (DYKDHDGDYKDHDIDYKDDDDK) was fused to the C-terminus of UNC93B1. TLR9, TLR7, and TLR3 were fused to HA (YPYDVPDYA) at the C-terminal end. TLR7 sequence was synthesized after codon optimization by Invitrogen’s GeneArt Gene Synthesis service as previously described^15^. TLR9 chimeras of the juxtamembrane and transmembrane regions were previously described^4^.

### Cells and tissue culture conditions

HEK293T cells were obtained from American Type Culture Collection (ATCC). GP2-293 packaging cell lines were obtained from Clontech. The above cell lines were cultured in DMEM complete media supplemented with 10% (vol/vol) FCS, L-glutamine, penicillin-streptomycin, sodium pyruvate, and HEPES (pH 7.2) (Invitrogen). RAW264 macrophage cell lines (ATCC) were cultured in RPMI 1640 (same supplements as above).

BMMs were differentiated for seven days in RPMI complete media (same supplements as above plus 0.00034% (vol/vol) beta-mercaptoethanol) and supplemented with M-CSF containing supernatant from 3T3-CSF cells.

To generate HEK293T Unc93b1^-/-^ cells, guide RNAs were designed and synthesized as gBlocks as previously described^24^ and then were subcloned into pUC19 (guide RNA: CTCACCTACGGCGTCTACC). Humanized Cas9-2xNLS-GFP was a gift from the Doudna laboratory, University of California, Berkeley, CA. HEK293T cells were transfected using Lipofectamine LTX with equal amounts of the guide RNA plasmid and Cas9 plasmid. Seven days post transfection cells were plated in a limiting-dilution to obtain single cells. Correct targeting was verified by PCR analysis and loss of response to TLR9 and TLR7 stimulation in an NFkB luciferase assay. Unc93b1^-/-^ RAW macrophages were generated with the Cas9(D10A)-GFP nickase (guide RNAs: 1) GGCGCTTGCGGCGGTAGTAGCGG, 2) CGGAGTGGTCAAGAACGTGCTGG, 3) TTCGGAATGCGCGGCTGCCGCGG, 4) AGTCCGCGGCTACCGCTACCTGG). Macrophages were transfected with cas9(D10A) and all four guide RNAs using Lipofectamine LTX and Plus reagent and single cell-sorted on cas9-GFP two days later. Correct targeting was verified by loss of response to TLR7 stimulation and sequencing of the targeted region after TOPO cloning.

### Retroviral transduction

For retroviral transduction of Raw macrophages, VSV-G-pseudotyped retrovirus was made in GP2-293 packaging cells (Clontech). GP2-293 cells were transfected with retroviral vectors and pVSV-G using Lipofectamine LTX reagent. 24h post-transfection, cells were incubated at 32°C. 48h post-transfection viral supernatant (with polybrene at final 5μg/ml) was used to infect target cells overnight at 32°C and protein expression was checked 48 hr later. Target cells were sorted on an Aria Fusion Beckman Coulter Sorter to match expression or drug-selected with Puromycin, starting 48h after transduction. Efficiency of drug selection was verified by equal mCherry expression of target cells.

For retroviral transduction of bone marrow derived macrophages, bone marrow was harvested and cultured in M-CSF containing RPMI for two days. Progenitor cells were transduced with viral supernatant (produced as above) on two successive days by spinfection for 90min at 32°C. 48h after the second transduction cells were put on Puromycin selection and cultured in M-CSF containing RPMI media until harvested on day 8.

### Luciferase assays

Activation of NF-κB in HEK293T cells was performed as previously described^8^. Briefly, transfections were performed in OptiMEM-I (Invitrogen) with LTX transfection reagent (Invitrogen) according to manufacturer’s guidelines. Cells were stimulated with CpG-B (200nM – 1μM), R848 (100-200ng/ml), or human IL-1b (20ng/ml) after 24h and lysed by passive lysis after an additional 12–16h. Luciferase activity was measured on a LMaxII-384 luminometer (Molecular Devices).

### Immunoprecipitation and western blot analysis

Cells or purified phagosomes were lysed in NP-40 buffer (50 mM Tris [pH 7.4], 150 mM NaCl, 1% NP-40, 5 mM EDTA and supplemented with EDTA-free complete protease inhibitor cocktail; Roche and 1mM PMSF). After incubation at 4°C on a rotator, lysates were cleared of insoluble material by centrifugation. For immunoprecipitations, lysates were incubated with anti-HA matrix, or anti-FLAG matrix (both pre-blocked with 1% BSA-PBS) overnight, and washed four times in lysis buffer the next day. Precipitated proteins were denatured in SDS-PAGE buffer at room temperature for 1h. Proteins were separated by SDS-PAGE (Bio-Rad TGX precast gels [Bio-Rad]) and transferred to Immobilon PVDF membranes (Millipore) in a Trans-Blot Turbo transfer system (Biorad). Membranes were probed with the indicated antibodies and developed using the Licor Odyssey Blot Imager. Relative band intensities were quantified using Fiji (ImageJ)^25^.

Streptavidin pull downs were performed on cells fed biotin-CpG-B for 4h, lysed in NP-40 buffer (same as above) and cleared of insoluble debris. Lysates were incubated for 2?h with streptavidin magnetic beads (pre-blocked with 1% BSA-PBS), rotated at 4?°C, and washed four times in lysis buffer. Precipitates were boiled in SDS buffer, separated by SDS–PAGE, and probed by anti-HA immunoblot. Cell lysis and co-immunoprecipitations for Myd88 pull downs were performed in the following buffer: 50mM Tris-HCl pH 7.4, 150mM NaCl, 10% glycerol, 1% NP-40 and supplemented with EDTA-free complete protease inhibitor cocktail (Roche), PhosSTOP (Roche) and 1mM PMSF. Lysates were incubated overnight with anti-Myd88 antibody, rotating at 4°C, and then Protein G agarose (pre-blocked with 1% BSA-PBS) was added for additional 2h. Beads were washed four times in lysis buffer, incubated in SDS buffer at room temperature for 1h, separated by SDS-PAGE, and probed for the indicated antibodies.

For visualizing disulfide bond formation in RAW macrophages, cells were lysed in buffer containing 10% DDM/CHS detergent for 2h at 4°C. After removing insoluble material, lysates were incubated with HA matrix for 2-4h and washed four times in buffer containing 0.25% DDM/CHS. Protein was eluted in lysis buffer containing 10% DDM/CHS and 300μg/ml HA peptide for 1h at room temperature. Eluates were divided in half and denatured in either reducing (+DTT) or non-reducing (-DTT) SDS buffer for 1h at room temperature.

### Flow cytometry

Cells were seeded into non-treated tissue culture 24-well plates or round-bottom 96-well plates. The next day cells were stimulated with the indicated TLR ligands. To measure TNFα production, BrefeldinA (BD GolgiPlug, BD Biosciences) was added to cells 30 min after stimulation, and cells were collected after an additional 5.5h. Cells were stained for intracellular TNF*a* with a Fixation & Permeabilization kit according to manufacturer’s instructions (eBioscience).

For measuring DNA uptake, cells were fed Cy3-fluorescent CpG-B for the indicated amounts of time (control cells for no uptake were pre-chilled and stimulated on ice). Cells were washed 3 times with ice-cold PBS, fixed in 1% PFA/2%FCS/PBS, and analyzed on an BD LSR Fortessa flow cytometer.

### Enzyme-linked immunosorbent assay (ELISA)

Cells were seeded into tissue culture-treated flat-bottom 96-well plates. The next day cells were stimulated with the indicated TLR ligands. For TNFa ELISAs NUNC Maxisorp plates were coated with anti-TNFa at 1.5μg/ml overnight at 4°C. Plates were then blocked with PBS + 1% BSA (w/v) at 37°C for 1h before cell supernatants diluted in PBS + 1% BSA (w/v) were added and incubated at room temperature for 2hrs. Secondary anti-TNFa biotin was used at 1μg/ml followed by Streptavidin-HRP. Plates were developed with 1mg/mL OPD in Citrate Buffer (PBS with 0.05M NaH2 PO4 and 0.02M Citric acid, pH 5.0) with HCl acid stop.

### Quantitative PCR

Cells were lysed in RNAzol (Molecular Research Center) and RNA was purified using the Direct-zol RNA MiniPrep Plus kit (Zymo Research) according to manufacturer’s instructions. RNA was treated with RQ1 RNase-free DNase (Promega) and concentrated using the RNA clean and concentrator-5 Kit (Zymo Research). cDNA was prepared from 100-500ng RNA with iScript cDNA synthesis kit, and quantitative PCR was performed with SYBR green on a StepOnePlus thermocycler (Applied Biosystems). Primers were synthesized by Integrated DNA Technologies. Primer sequences (PrimerBank): Ifnb1 F: AGCTCCAAGAAAGGACGAACA, Ifnb1 R: GCCCTGTAGGTGAGGTTGAT; Tnfa F: CAGGCGGTGCCTATGTCTC, Tnfa R: CGATCACCCCGAAGTTCAGTAG; Gapdh F: AGGTCGGTGTGAACGGATTTG, Gapdh R: GGGGTCGTTGATGGCAACA

### Microscopy

Coverslips (High-performance, 18×18mm, thickness no. 1 ½ Zeiss) were acid-washed in 3M HCl, washed extensively in water, dipped in 70% EtOH and allowed to air-dry. Cells were plated onto coverslips and allowed to settle overnight. The next day, cells were incubated with Cy3-CpG-A for 2hrs at 37°C. Coverslips were washed with PBS, fixed with 4% PFA/PBS for 15min, and permeabilized with 0.5% saponin/PBS for 5min. To quench PFA autofluorescence coverslips were treated with sodium borohydride/0.1% saponin/PBS for 10min. After washing 3x with PBS, cells were blocked in 1% BSA/0.1% saponin/PBS for 1h. Slides were stained in blocking buffer with anti-HA, and anti-LAMP1 (see antibodies above), washed with PBS and incubated for 45min with secondary antibodies. Cells were washed 3x in PBS and mounted in VectaShield Hard Set without Dapi. Cells were imaged on a Zeiss Elyra PS.1 with a 100x/1.46 oil immersion objective in Immersol 518F / 30°C (Zeiss). Z-Sections were acquired, with three grid rotations at each Z-position. The resulting dataset was SIM processed and Channel Aligned using Zeiss default settings in Zen. The completed super-resolution Z-Series was visualized and analyzed using Fiji (ImageJ). To compare the degree of colocalization of two proteins a single section from the middle of the Z-Series was selected and analyzed using a customized pipeline for object-based colocalization in Cell Profiler^26^. Briefly, primary objects (Unc93b1 vs Lamp1, or CpG-A vs TLR9) were identified and related to each other to determine the degree of overlap between objects. Data are expressed as % of object 1 colocalized with object 2. For some colocalization experiments, pixel intensities of two different fluorophores were measured by using the Plot Profile tool in Fiji to create a plot of intensity values along a line scan in the image.

### Phagosome isolation

Cells in a confluent 15cm dish were incubated with ~10^8^ 1?μm magnetic beads (Polysciences) for 4hrs. After rigorous washing in PBS, cells were scraped into 10?ml sucrose homogenization buffer (SHB: 250?μM sucrose, 3?mM imidazole pH?7.4) and pelleted by centrifugation. Cells were resuspended in 2?ml SHB plus protease inhibitor cocktail with EDTA (Roche) and 1mM PMSF and disrupted by 25 strokes in a steel dounce homogenizer. The disrupted cells were gently rocked for 10?min on ice to free endosomes. Beads were collected with a magnet (Dynal) and washed 4x with SHB plus protease inhibitor. After the final wash, phagosome preparations were denatured in 2x SDS buffer at room temperature for 1h and analyzed by western blot.

### Cell fractionation by sucrose density-centrifugation

Cells in four confluent 15cm dishes were washed in ice-cold PBS, scraped in 10ml sucrose homogenization buffer (SHB: 250?μM sucrose, 3?mM imidazole pH?7.4) and pelleted by centrifugation. Cells were resuspended in 2?ml SHB plus protease inhibitor cocktail with EDTA (Roche) and 1mM PMSF and disrupted by 25 strokes in a steel dounce homogenizer. The disrupted cells were centrifuged for 10min at 1000g to remove nuclei. Supernatants were loaded onto continuous sucrose gradients (percent iodixanol: 0, 10, 20, 30) and ultracentrifuged in an SW41 rotor at 25800rpm for 2hrs (Optima L-90K Ultracentrifuge, Beckman Coulter). 22 fractions of 420μl were collected from top to bottom. 100μl of each fraction were denatured in SDS buffer for western blot analysis. For immunoprecipitations, three fractions corresponding to ER or endosomes were combined and lysed for 1h after addition of protease inhibitor cocktail and NP-40 to a final concentration of 1%. Coimmunoprecipitation with anti-HA matrix was performed as described above.

### Quantification and Statistical Analysis

Statistical parameters, including the exact value of n and statistical significance, are reported in the Figures and Figure Legends. Representative images have been repeated at least three times, unless otherwise stated in the figure legend. Data is judged to be statistically significant when p < 0.05 by two-tailed Student’s t-test. To compare the means of different groups, a one-way ANOVA followed by a Tukey’s posttest was used. To compare means of different groups that have been split on two independent variables, a two-way ANOVA followed by a Bonferroni posttest was used. In figures, asterisks denote statistical significance (*, p < 0.05; **, p < 0.01; ***, p < 0.001). Statistical analysis was performed in GraphPad PRISM 7 (Graph Pad *Software* Inc.).

## Acknowledgments

We thank members of the Barton and Vance Lab for helpful discussions and critical reading of the manuscript. We thank Baobin Li and Stephen Brohawn for supplying the DDM/CHS detergent and for technical advice. We thank Hector Nolla and Alma Valeros for assistance with cell sorting at the Flow Cytometry Facility of the Cancer Research Laboratory at UC Berkeley. We thank Steven Ruzin and Denise Schichnes for assistance with microscopy on the Zeiss Elyra PS.1 at the Biological Imaging Center at UC Berkeley. This work was supported by the NIH (AI072429, AI105184 and AI063302 to G.M.B.) and by the Lupus Research Institute (Distinguished Innovator Award to G.M.B.). O.M. was supported by an Erwin Schrödinger (J 3415-B22) and CRI Irvington postdoctoral fellowship. B.J.W. was supported by a summer undergraduate research fellowship from UC Berkeley. B.L. was supported by the UC Berkeley Tang Distinguished Scholars Program. Research reported in this publication was supported in part by the National Institutes of Health S10 program under award number 1S10OD018136-01.

## Author Contributions

O.M. and G.M.B designed experiments. O.M., B.J.W., and E.V.D. performed experiments and analyzed the data. O.M. and B.L. performed the initial alanine mutagenesis screen. O.M. wrote the manuscript. G.M.B. revised and edited the manuscript.

## Supplementary Materials for

**Fig. S1.**
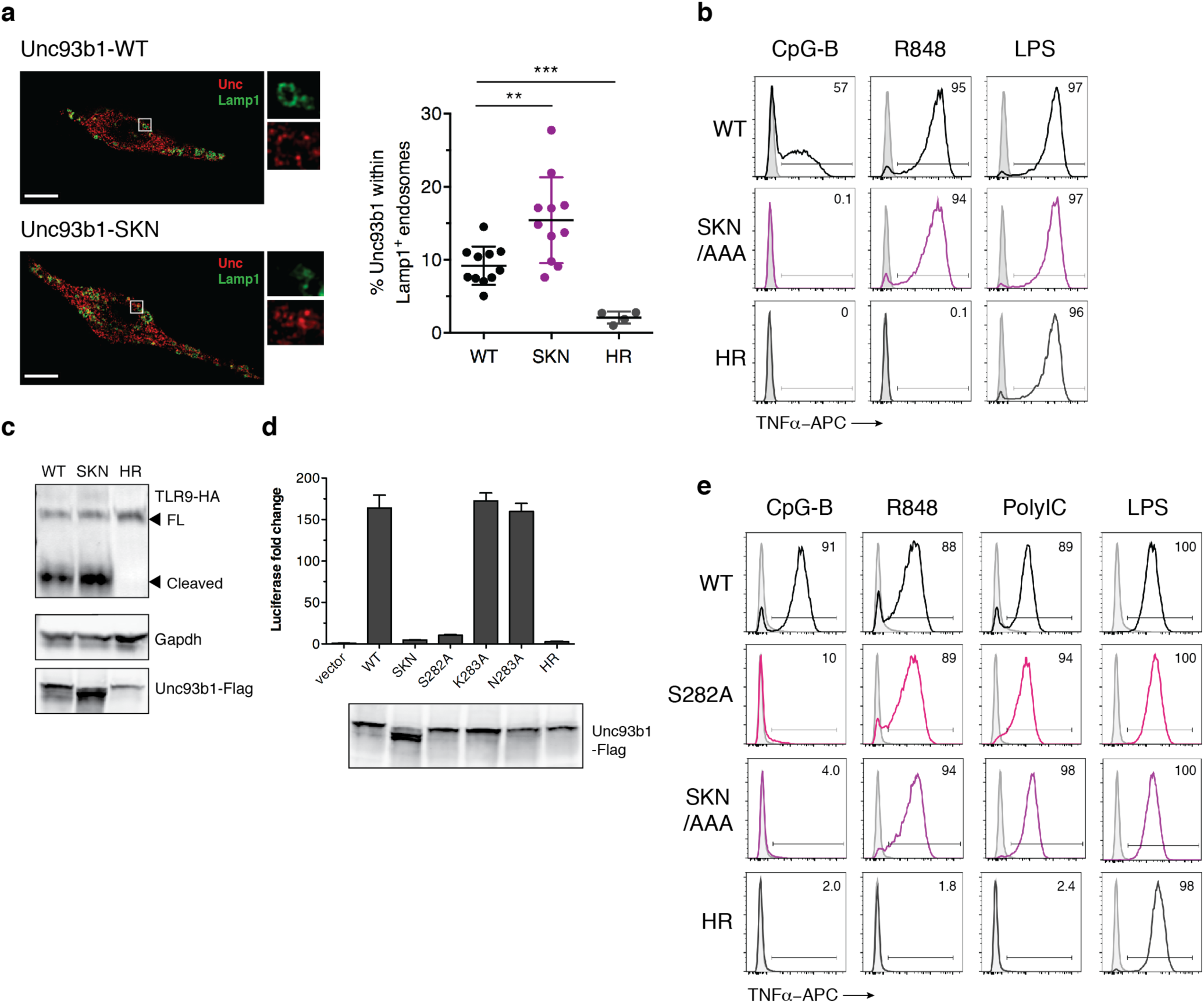
A luminal Unc93b1 mutation results in defective TLR9 signaling despite normal trafficking. (**a**) Colocalization of Unc93b1 and Lamp1 in RAW macrophages expressing the indicated Unc93b1 alleles using superresolution structured illumination microscopy. Shown are representative Unc93b1^WT^ and Unc93b1^SKN^ cells: Unc93b1 (red) and Lamp1 (green). Boxed areas are magnified. The plot shows quantification of the percentage of total Unc93b1 within Lamp1^+^ endosomes. Each dot represents an individual cell imaged in a single experiment. Scale bars: 10μm. (**b**) Unc93b1^SKN^ selectively impairs TLR9 signaling in primary macrophages. Representative flow cytometry analysis showing percent TNFα positive cells, measured by intracellular cytokine staining, of BMMs from *Tlr9*^HA:GFP^ *Unc93b1*^-/-^ mice expressing the indicated Unc93b1 alleles (WT, SKN, and non-functional HR mutant) after stimulation with CpG-B (1μM) for TLR9, R848 (50ng/ml) for TLR7, and LPS (10ng/ml) for TLR4. (**c**) TLR9 trafficking is normal in Unc93b1^SKN^-expressing BMMs. Immunoblot of TLR9 from lysates of the same BMMs shown in (**b**); FL: full-length. (**d**) Unc93b1^S282A^ is sufficient for the TLR9 signaling defect.NF?B activation was measured by luciferase assay in HEK293T cells transiently transfected with TLR9 and the indicated Unc93b1 alleles and stimulated with CpG-B (1μM) for 16hrs. Data is normalized to Unc93b1-independent hIL-1b responses and expressed as luciferase fold change over unstimulated controls (n=4). Blot below shows Unc93b1 expression levels. (**e**) Unc93b1^S282A^ recapitulates the Unc93b1^SKN^-mediated TLR9 signaling defect in RAW macrophages. Representative flow cytometry analysis showing percent TNFα positive cells, measured by intracellular cytokine staining, of RAW macrophage lines expressing the indicated Unc93b1 alleles after stimulation with CpG-B (1μM) for TLR9, R848 (100ng/ml) for TLR7, PolyIC (100ng/ml) for TLR3, and LPS (10ng/ml) for TLR4. RAW macrophages used here were retrovirally transduced to express TLR3-HA. All data are mean ± SD; *P < 0.05, **P < 0.01, ***P < 0.001 by unpaired Student’s t-test. The data are representative of at least two independent experiments, unless otherwise noted.

**Fig. S2.**
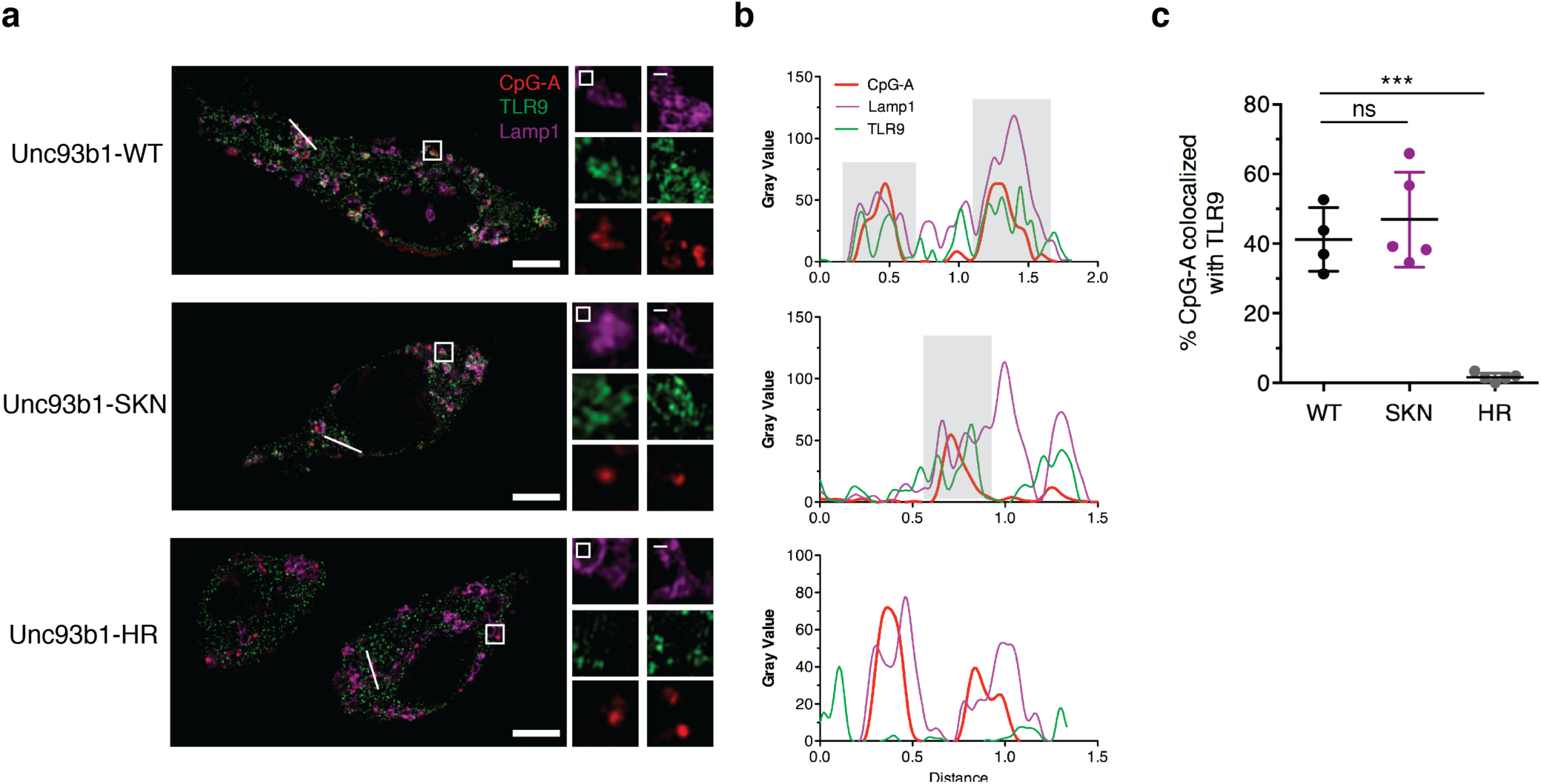
Unc93b1^S282A^ does not affect the DNA delivery to TLR9-containing endosomes. Colocalization of CpG-A, TLR9, and Lamp1 was determined in RAW macrophage lines expressing the indicated Unc93b1 alleles after incubation with Cy3-labeled CpG-A (1μM) for 2hrs using superresolution structured illumination microscopy. (**a**) Shown are representative cells: TLR9 (green), Lamp1 (magenta), and CpG-A (red). Boxed areas and areas containing white lines are magnified. (**b**) The histograms display fluorescent intensity plots of pixels along the white lines. Shaded areas highlight regions of colocalization of CpG-B, TLR9, and Lamp1. (**c**) The plot shows quantification of the percentage of CpG-A colocalized with TLR9. Each dot represents an individual cell imaged in a single experiment. Scale bars: 5μm. Data are mean ± SD; *P < 0.05, **P < 0.01, ***P < 0.001 by unpaired Student’s t-test.

**Fig. S3.**
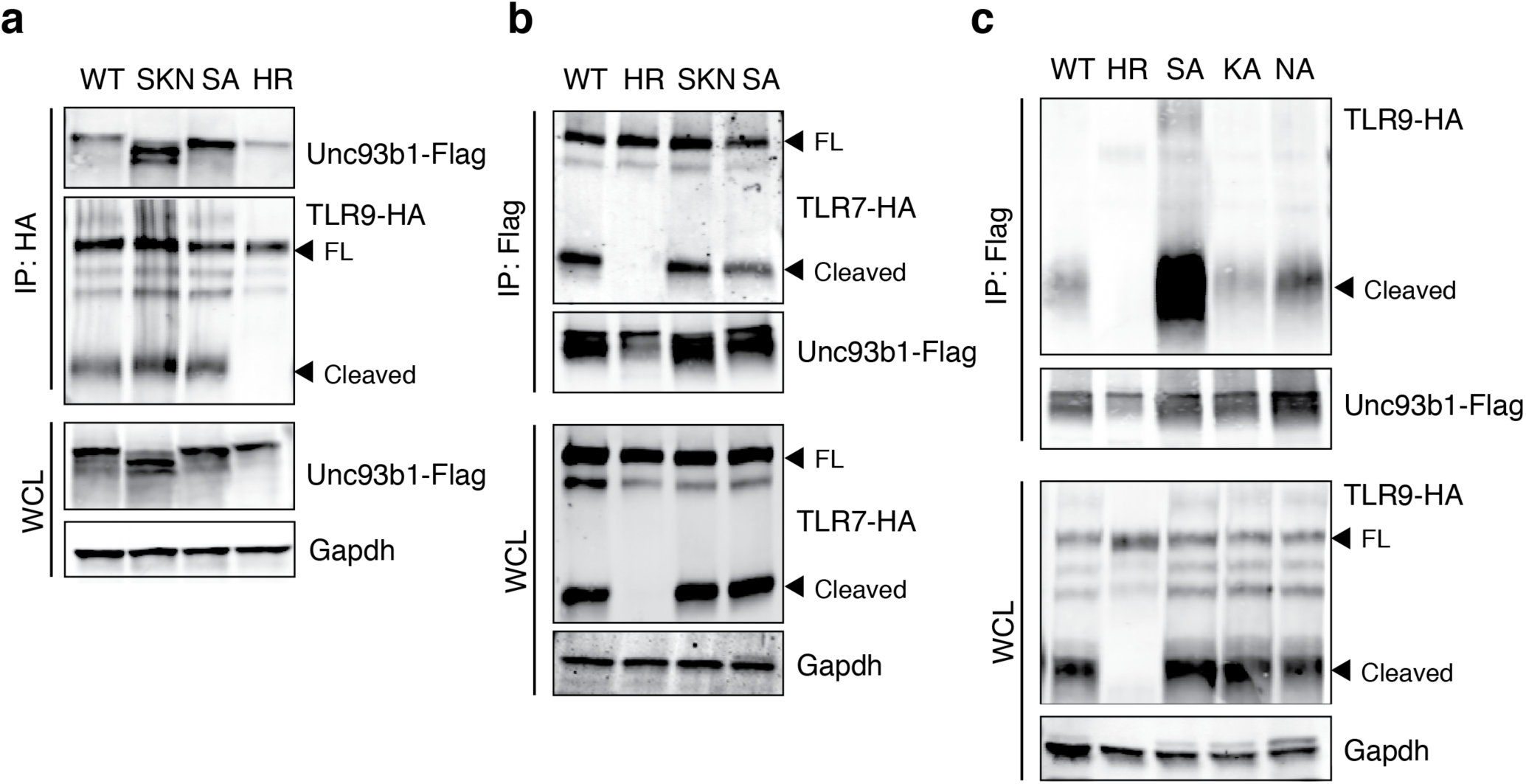
The stronger interaction is specific to Unc93b1^S282A^ and TLR9. (**a**) Unc93b1^SKN^ and Unc93b1^S282A^ bind stronger to TLR9. HA immunoprecipitation of TLR9 from RAW macrophage lines, expressing the indicated Unc93b1 alleles, followed by immunoblot of Unc93b1. Input levels of Unc93b1 in whole cell lysates (WCL) are also shown. (**b**) Unc93b1^SKN^ and Unc93b1^S282A^ do not affect the interaction with TLR7. Flag immunoprecipitation of Unc93b1 from RAW macrophage lines, expressing TLR7-HA and the indicated Unc93b1 alleles, followed by immunoblot of TLR7. Input levels of TLR7 in whole cell lysates (WCL) are also shown. (**c**) Only Unc93b1^S282A^, but not Unc93b1^K283A^ or Unc93b1^N284A^, bind stronger to TLR9. Flag immunoprecipitation of Unc93b1 from RAW macrophage lines, expressing the indicated Unc93b1 alleles, followed by immunoblot of TLR9. Input levels of TLR9 in whole cell lysates (WCL) are also shown; FL: full-length. The data are representative of at least two independent experiments, unless otherwise noted.

**Fig. S4.**
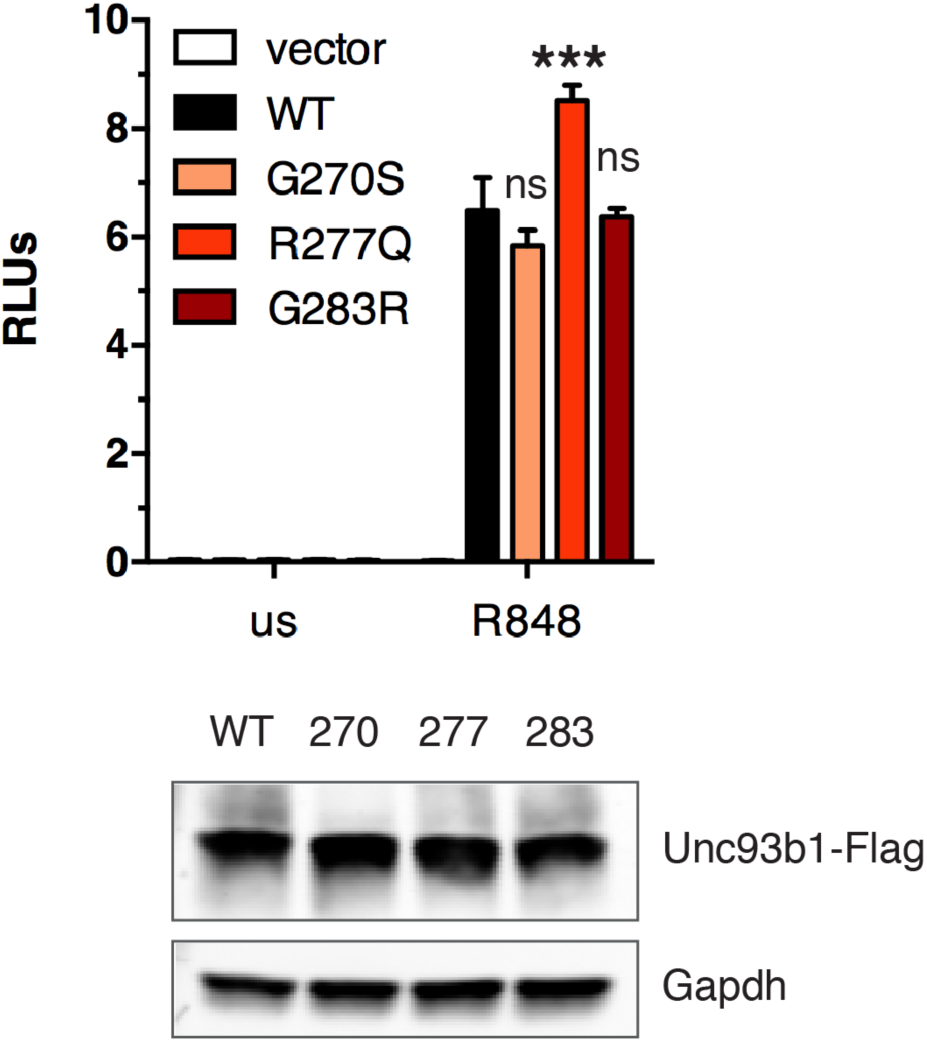
Human Unc93b1 variants do not reduce TLR7 signaling. NFκB activation was measured by luciferase assay in HEK293T cells transiently transfected with TLR7 and the indicated human Unc93b1 variants and stimulated with R848 (250ng/ml) for 16hrs. Data are normalized to Renilla expression and expressed as relative luciferase units (RLUs) (n=3, representative of three independent experiments). Expression levels of human Unc93b1 variants are shown below. All data are mean ± SD; *P < 0.05, **P < 0.01, ***P < 0.001 by one-way ANOVA followed by a Tukey’s posttest. On-way ANOVA results: F(3/8)=29.70, *p*=0.0001

**Fig. S5.**
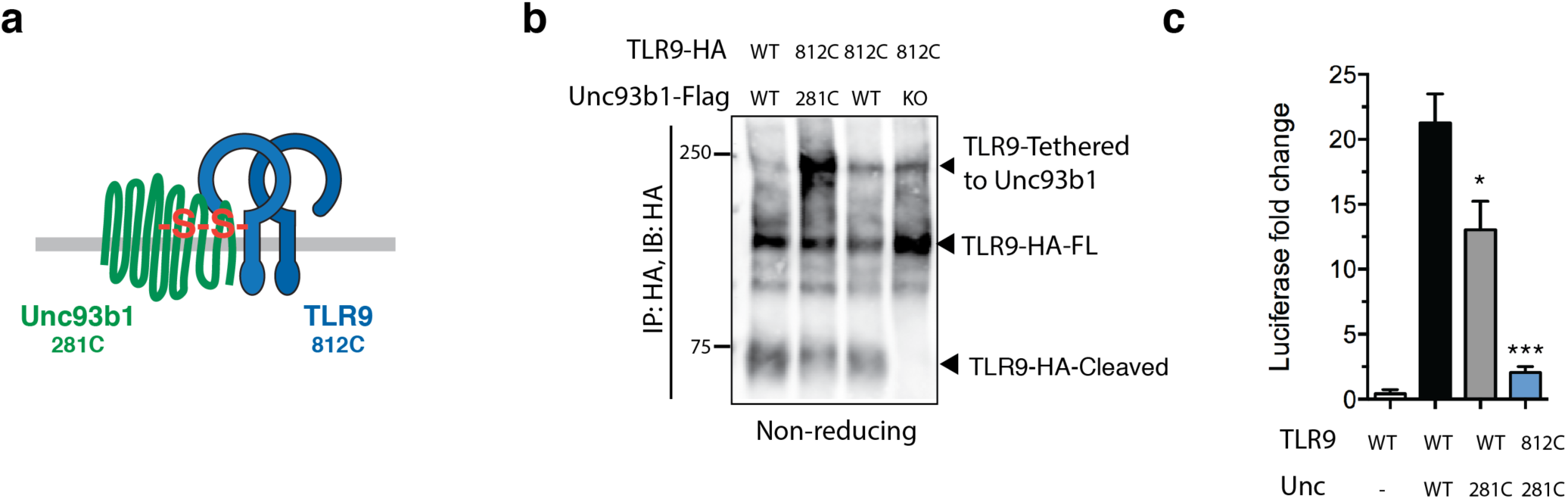
Tethering of TLR9 and Unc93b1. (**a**) Schematic of the disulfidebond-tethered Unc93b1-TLR9 complex. The engineered cysteine position in either protein is indicated. (**b**) Unc93b1-tethered TLR9 is unable to signal. HA immunoblot under non-reducing conditions after HA immunoprecipitation of TLR9 from RAW macrophage lines expressing the indicated TLR9 and Unc93b1 cysteine mutants. The high molecular weight band indicates disulfide-bond formation between Unc93b1 and TLR9. Representative blot out of two independent experiments. (**c**) NF?B luciferase assay in HEK293T cells expressing the indicated cysteine mutant combinations and stimulated with CpG-B (1μM) for 16hrs. Data are normalized to Renilla expression and expressed as luciferase fold change over unstimulated controls (n=3, representative of three independent experiments).

